# Integrative analysis across metagenomic taxonomic classifiers: A case study of the gut microbiome in aging and longevity

**DOI:** 10.1101/2025.05.14.654104

**Authors:** Tanya T. Karagiannis, Ye Chen, Sarah Bald, Albert Tai, Sofiya Milman, Stacy L. Andersen, Thomas T. Perls, Daniel Segrè, Paola Sebastiani, Meghan I. Short

## Abstract

Despite calls for the development of consensus methods, most analyses of shotgun metagenomics data for microbiome studies use a single taxonomic classifier. In this study, we compare inferences from two broadly used classifiers, MetaPhlAn4 (marker-gene-based) and Kraken2 (k-mer-based), applied to stool metagenomic samples from participants in the Integrative Longevity Omics study to measure associations of taxonomic diversity and relative abundance with age, replicating analyses in an independent cohort. We also introduce consensus and meta-analytic approaches to compare and integrate results from multiple classifiers. While many results are consistent across the two classifiers, we find classifier-specific inferences that would be lost when using one classifier alone. When using a correlated meta-analysis approach across classifiers, differential abundance analysis captures more age-associated taxa, including 17 taxa robustly age-associated across cohorts. This study emphasizes the value of employing multiple classifiers and recommends novel approaches that facilitate the integration of results from multiple methodologies.

## INTRODUCTION

High-throughput sequencing approaches, such as shotgun metagenomics, have greatly increased our ability to investigate changes in microbial communities of the human gut.^1^ A variety of taxonomic classification methods are available for processing shotgun metagenomics sequencing data into taxonomic profiles of microbial communities. Taxonomic profilers identify taxa and their relative abundances in each sample based on the classification of sequencing reads against a reference database. The methods underlying these classification tools vary and include marker-gene-based methods (e.g., MetaPhlAn^2^, mOTUs^3^), k-mer-based approaches (e.g. Kraken^4,5^, Bracken^6,7^, Centrifuge^8^), and protein-based methods (e.g. Kaiju^9^, DIAMOND^10^), each with different strengths.

Two popular methods, MetaPhlAn and Kraken, have been consistently used in metagenomics studies, including in the context of aging. MetaPhlAn performs classification of sequencing reads by aligning reads against a curated database of marker genes specific to taxonomic groups, with species relative abundances estimated based on clade-specific coverage.^2,11^ Although this method does not use all the sequencing data, MetaPhlAn has been shown to have high specificity and high coverage of the human gut microbiome.^2,12^ Kraken performs k-mer based classification by mapping individual reads to taxa via k-mer “voting” methods.^4^ More specifically, Kraken maps reads to the lowest taxonomic group in the taxonomic hierarchy that shares that k-mer to infer taxonomic abundance, with estimation of relative abundances at a specific level performed by Bracken.^4,6^ Although the Kraken/Bracken approach has greater sensitivity than marker-based methods and uses all the sequencing data, it has been shown to be prone to false positives.^12-15^ These differences between classification approaches can greatly impact the identification of taxa, their estimated relative abundances, and downstream analyses.^11,16^

Because of the differences and complementary strengths of various taxonomic classification approaches, previous work has suggested that consensus-based methods may be desirable for robust microbiome analysis.^14,17,18^ However, there is little guidance or examples of this in practice. Tools, such as MetaMeta^19^, WEVOTE^20^, FlexTaxD^21^, and a recently developed *merging strategy* using a weighted voting approach^22^ have provided avenues for analysis of metagenomics data via multiple taxonomic classification methods. These integrative approaches use a variety of methods to combine classifier-specific profiles and improve accuracy of taxonomic identification. However, these methods are limited by the range of classification tools supported—e.g., MetaMeta cannot support MetaPhlAn, FlexTaxD can only support k-mer based classifiers, *merging strategy* was tested on a limited number of profilers excluding marker-gene based methods—and some methods such as MetaMeta or WEVOTE are not maintained by the developers. Additionally, these approaches all perform integration at the taxonomic classification level, creating a single combined feature table for downstream analysis. Combined classification in this way precludes comparison of taxonomic profiles and downstream findings from each profiler method. Finally, despite the availability of some tools and the potential benefits of analysis with multiple classifiers, most studies continue to rely on a single taxonomic profiler.^17,23,24^

Motivated by the interest in comparing analysis results from different classifiers and leveraging their complementary strengths, we used two taxonomic profiling approaches in parallel (Kraken2 and MetaPhlAn4) to discover diversity trends and individual taxa associated with age in two studies of extreme human longevity. We performed shotgun metagenomics sequencing of the gut microbiome from individuals enrolled in the Integrative Longevity Omics Study (ILO), a new cohort study of centenarians in North America and their offspring. In addition, we acquired publicly available metagenomic sequence data from a cohort of Han Chinese individuals as a replication cohort.^25^ We present this analysis as a case study to show the value of using complementary approaches to analyze metagenomics data and introduce methods, including a novel correlated meta-analysis approach, that can help integrate results across taxonomic classifiers and provide comprehensive analysis of the data.

## RESULTS

### Two popular taxonomic classifiers detect different taxonomic profiles

We generated shotgun metagenomics sequencing data of the gut microbiome from 220 participants of the Integrative Longevity Omics (ILO) study (59-107 years), who were of North American/European descent and included 78 centenarians (100-107 years) and 142 of their biological offspring (59-99 years) (**Figure 1, see Supplementary Methods**). We also obtained a shotgun metagenomics sequencing dataset of the gut microbiome from 348 individuals of Han Chinese descent (50-105 years), with 116 considered to be of advanced age (90-105 years) including 13 centenarians (100-105 years), and 232 of their offspring (50-79 years)^25^ (**Figure 1**). We processed both raw sequence data sets using KneadData^26^, and performed taxonomic classification using 1) MetaPhlAn4^2^ and 2) Kraken2^4^ followed by Bracken.^6^

**Figure 1.**
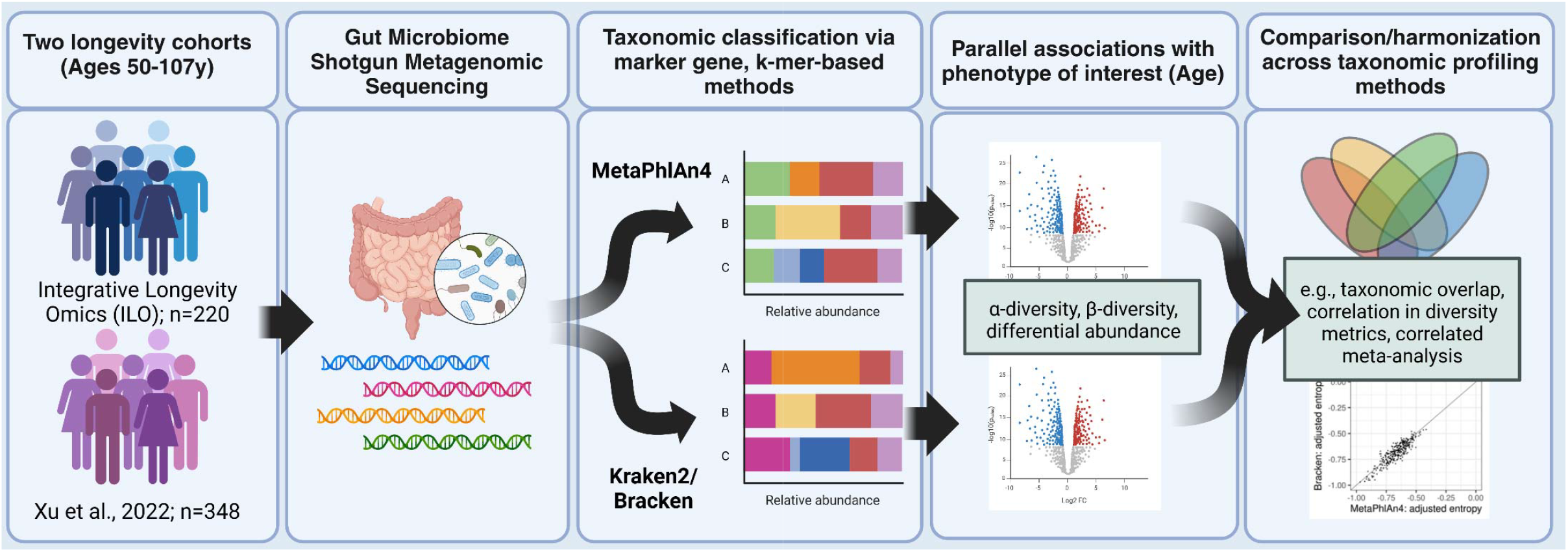
Overview of analysis pipeline for shotgun metagenomics analysis using complementary taxonomic classification approaches. Shotgun metagenomics sequencing data of the gut microbiome was generated from individuals of the Integrative Longevity Omics Study (ILO), and acquired from a cohort of Han Chinese individuals as a replication cohort. We used two complementary taxonomic profiling approaches in parallel (Kraken2 and MetaPhlAn4) to discover taxa associated with age from the two studies of extreme human longevity. We measured associations of alpha and beta diversity with age, as well as differential abundance of taxa with age, using profiles generated by both classifiers in both cohorts. We used taxonomic overlap, correlation, Procrustes analysis, voting methods, and a correlated meta-analysis approach recently developed in our research group to measure agreement and integrate downstream results across classifiers.

There were differences in species numbers between the databases used by MetaPhlAn4 (20789 species) and Kraken2/Bracken (23127 species), with relatively minimal overlap (6292 species in common, or 27-30% of species in each database) (**Table 1**). After taxonomic classification and filtering procedures were applied to the cohort datasets, Kraken2/Bracken identified more species (ILO:1044, Xu:1504) than MetaPhlAn4 (ILO:787, Xu:898). The within-cohort overlap in the species identified by the two profilers was low (ILO:335 (22.4%), Xu:442 (21.3%)), and greater overlap observed within the same profiler across cohort (MetaPhlAn4:626 (59.1%), Kraken2/Bracken:846 (49.7%)). We observed similar trends across most taxonomic levels (**Table S1**).

**Table 1.**
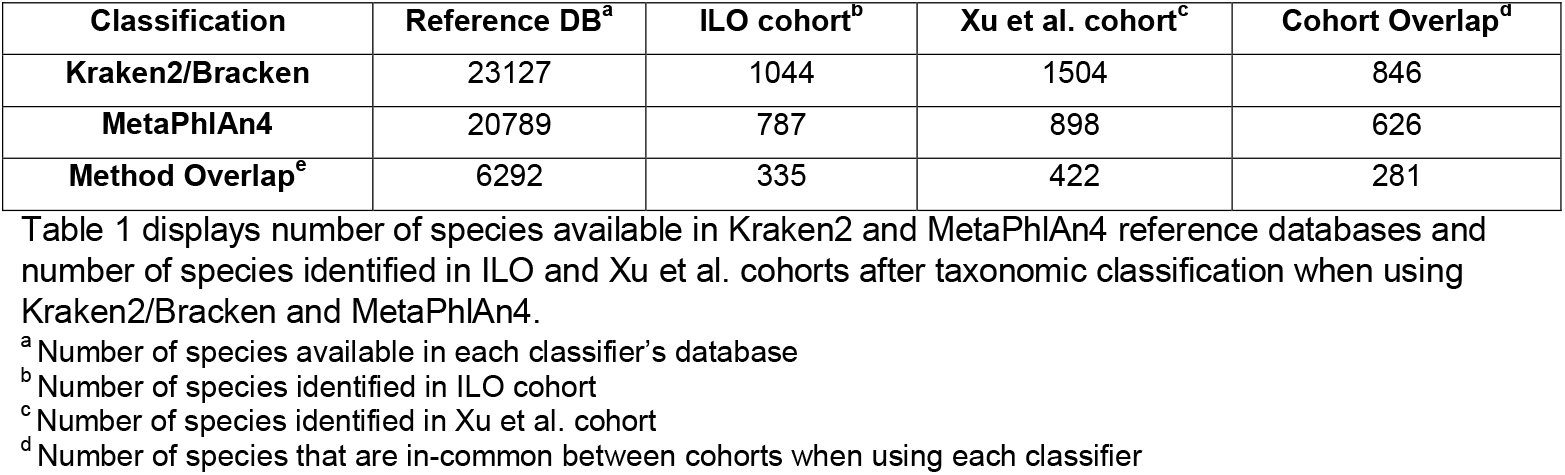
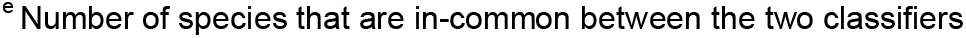
Number of species identified by MetaPhlAn4 and Kraken2/Bracken.

| Classification | Reference DB<sup>a</sup> | ILO cohort<sup>b</sup> | Xu et al. cohort<sup>c</sup> | Cohort Overlap<sup>d</sup> |
| --- | --- | --- | --- | --- |
| Kraken2/Bracken | 23127 | 1044 | 1504 | 846 |
| MetaPhlAn4 | 20789 | 787 | 898 | 626 |
| Method Overlap<sup>e</sup> | 6292 | 335 | 422 | 281 |

Table 1 displays number of species available in Kraken2 and MetaPhlAn4 reference databases and number of species identified in ILO and Xu et al. cohorts after taxonomic classification when using Kraken2/Bracken and MetaPhlAn4.
<sup>a</sup> Number of species available in each classifier's database <sup>b</sup> Number of species identified in ILO cohort <sup>c</sup> Number of species identified in Xu et al. cohort <sup>d</sup> Number of species that are in-common between cohorts when using each classifier
<sup>e</sup> Number of species that are in-common between the two classifiers

Since we found differences in the species identified across classifiers, we next investigated if the two methods differed by their quantification of relative abundances among species identified by both classifiers. The taxonomic profiles generated by Bracken and MetaPhlAn4 varied substantially based on the composition of in-common species identified by both classifiers.

These shared species comprised the bulk of the sample profiles when using Bracken, representing an average of 88.1-90.0% of the total relative abundance in sample profiles of both cohorts. When using MetaPhlAn4, shared species comprised a smaller proportion of the total, making up, on average, 68.3-68.7% of the total relative abundance in the sample profiles of both cohorts (**Figure S2A-B**). However, the most abundant species in each profile were similar across classifier methods (**Figure S2C-D**), suggesting that the dominant taxa are similar regardless of the classifier used, and differences may result primarily from differential identification of lower abundance taxa. In addition, we investigated the unique taxa in each cohort dataset that did not overlap between taxonomic classifiers (**Table 2, Table S1**). Of the species found only by Kraken2/Bracken (709-1082 species), 52.7-61.2% were present in the MetaPhlAn4 database -- i.e., MetaPhlAn4 had the ability to identify those species but did not. In contrast, of the species found only by MetaPhlAn4 in each cohort dataset (452-476 species), only 1.6-2.3% were also present in the Kraken2/Bracken database -- i.e., Kraken2/Bracken had the ability to identify those species but did not. This suggests that Kraken2/Bracken has higher sensitivity than MetaPhlAn4 and/or potentially identifies more false positives, as has been suggested previously.^12,15^ We observed similar trends across most taxonomic levels (**Table S1**).

**Table 2.** Number of species identified by one classifier and their availability in the other classifier’s database.

| Classification | ILO cohort |  |  | Xu et al. cohort |  |  |
| --- | --- | --- | --- | --- | --- | --- |
|  | Unique taxa<sup>a</sup> | In Other DB<sup>b</sup> | Not in Other DB<sup>c</sup> | Unique taxa<sup>a</sup> | In Other DB<sup>b</sup> | Not in Other DB<sup>c</sup> |
| Kraken2/Bracken | 709 | 434<br>(61.21%) | 275<br>(38.79%) | 1082 | 570<br>(52.68%) | 512<br>(47.32%) |
| MetaPhlAn4 | 452 | 7<br>(1.55%) | 445<br>(98.45%) | 476 | 11<br>(2.31%) | 465<br>(97.69%) |

Table 2 displays number of species uniquely identified by one classifier method and the number of those species that are present or not present in the other classifier's database.
<sup>a</sup> Number of species uniquely identified by one classifier method
<sup>b</sup> Number of unique species identified by one classifier method that are also present in the other classifier's database
<sup>c</sup> Number of unique species identified by one classifier method that are not present in the other classifier's database

### Age associations with alpha diversity vary based on taxonomic level

We calculated a normalized alpha diversity of the taxonomic relative abundances at each taxonomic level using both classifiers (**Table S2, Supplementary Methods**). In both ILO and Xu et al. cohorts, we observed a significant increase of normalized alpha diversity with age at the phylum level with both classifiers (ILO MetaPhlAn4: slope=0.00103, p=0.00232; ILO Bracken: slope=0.00124, p=0.000314; Xu MetaPhlAn4: slope=0.00159, p=7.6e-8; Xu Bracken: slope=0.00084, p=0.00317) (**Figure 2A-B**). This trend in diversity with age persisted at the class and order levels of the taxonomy (**Figure S3A-B**). At the genus and species level, the increasing trend of normalized alpha diversity with age was not observed (**Figure 2A-B**). The results varied across classifier and cohort, with two cases at the species level showing significance although having flatter trends (ILO Bracken: slope=0.000655, p=0.0384; Xu MetaPhlAn4: slope=0.000681, p=0.0161). The significance of the slope observed when using Bracken in the ILO cohort may be influenced by outlier sample profiles with particularly low diversity (**Figure 2A**).

**Figure 2.**
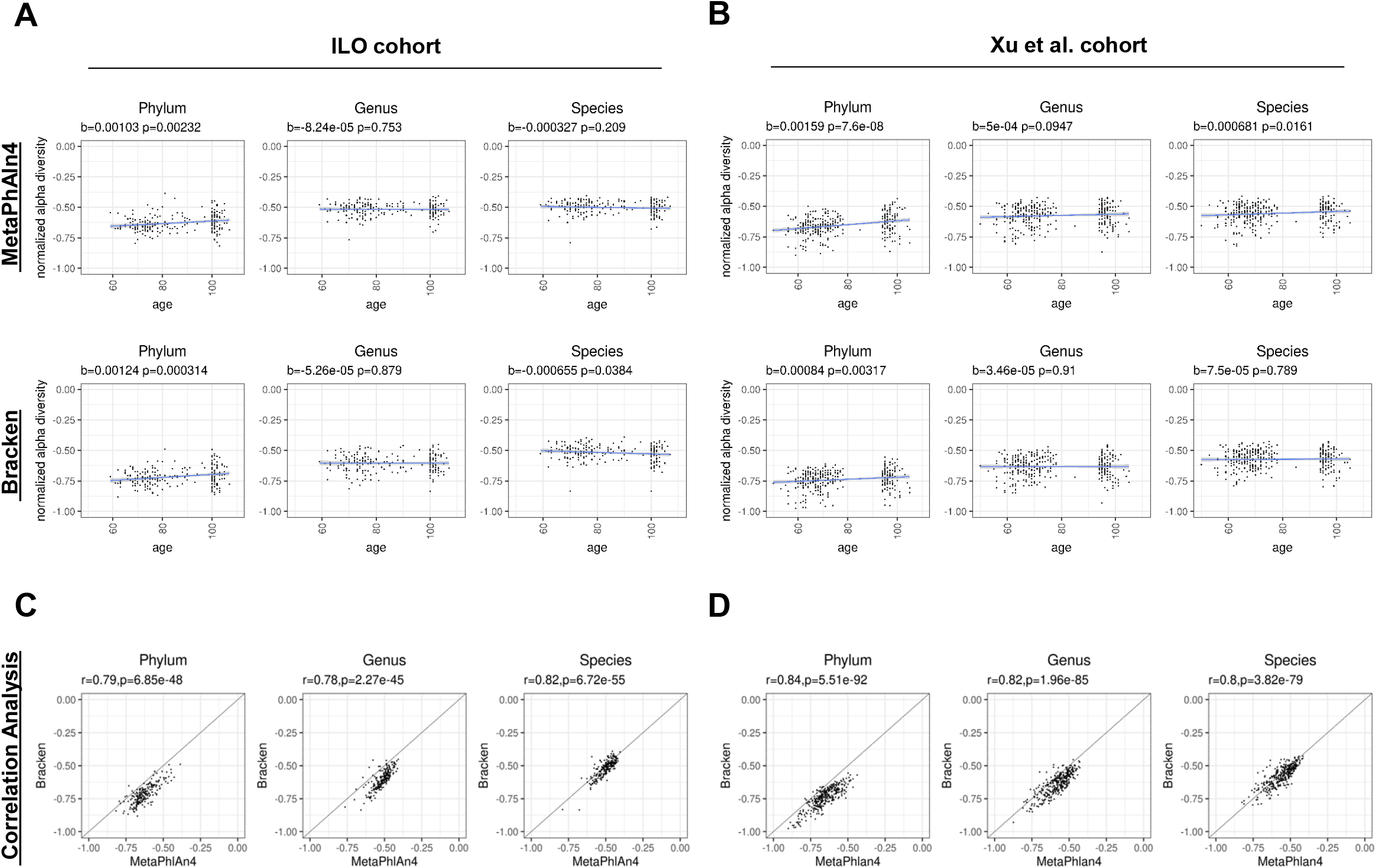
Alpha diversity displays similar changes with age at higher taxonomic levels and varies at lower taxonomic levels across classification approaches. **(A-B)** Scatterplots of the normalized alpha diversity score for each sample with age, comparing across phylum, genus, and species taxonomic levels within cohorts and between cohorts and classification methods. We employed linear regression models to evaluate the association with age, with a significance threshold set at 0.05. **(C-D)** Scatterplots comparing the normalized alpha diversity scores of samples based on classification method within each cohort. We employed the Pearson correlation analysis to evaluate differences in the sample normalized alpha diversity scores between methods, with significance threshold set at 0.05.

Normalized alpha diversities generated with MetaPhlAn4 and Kraken2/Bracken were highly correlated (Pearson correlation r>= 0.78, p<0.001 for all taxonomic levels) (**Figure 2C-D, Figure S3C-D**)(**Table 3**). The results were similar when restricted to in-common taxa identified by both classifier methods (**Figure S4**). Collectively, the normalized alpha diversity results suggest that both taxonomic classifiers capture consistent age associations in two cohorts at higher taxonomic levels, with inconsistent results at lower taxonomic levels.

**Table 3.** Correlation of diversity measures between classifiers within cohort.

| Cohort | Diversity Analysis<sup>a</sup> | Correlation Analysis<sup>b</sup> | Phylum<sup>c</sup> | Class<sup>d</sup> | Order<sup>e</sup> | Family<sup>f</sup> | Genus<sup>g</sup> | Species<sup>h</sup> |
| --- | --- | --- | --- | --- | --- | --- | --- | --- |
| ILO | Alpha | Pearson | r=0.79<br>p=6.85e-48 | r=0.79<br>p=2.67e-47 | r=0.79<br>p=3.09e-47 | r=0.76<br>p=4.14e-42 | r=0.78<br>p=2.27e-45 | r=0.82<br>p=6.72e-55 |
|  | Beta | Mantel | r=0.85<br>p≤0.001 | r=0.88<br>p≤0.001 | r=0.88<br>p≤0.001 | r=0.89<br>p≤0.001 | r=0.91<br>p≤0.001 | r=0.89<br>p≤0.001 |
|  |  | Procrustes | r=0.85<br>p≤0.001 | r=0.86<br>p≤0.001 | r=0.87<br>p≤0.001 | r=0.84<br>p≤0.001 | r=0.85<br>p≤0.001 | r=0.87<br>p≤0.001 |
| Xu | Alpha | Pearson | r=0.84<br>p=5.51e-92 | r=0.86<br>p=1.22e-100 | r=0.86<br>p=7.14e-101 | r=0.81<br>p=1.2e-79 | r=0.82<br>p=1.96e-85 | r=0.80<br>p=3.82e-79 |
|  | Beta | Mantel | r=0.78<br>p≤0.001 | r=0.79<br>p≤0.001 | r=0.79<br>p≤0.001 | r=0.83<br>p≤0.001 | r=0.86<br>p≤0.001 | r=0.86<br>p≤0.001 |
|  |  | Procrustes | r=0.83<br>p≤0.001 | r=0.83<br>p≤0.001 | r=0.84<br>p≤0.001 | r=0.83<br>p≤0.001 | r=0.84<br>p≤0.001 | r=0.86<br>p≤0.001 |

<sup>a</sup> Diversity analyses performed (Alpha or Beta diversity)
<sup>b</sup> Correlation analysis test performed to compare diversity measures between methods
<sup>c-h</sup> Taxonomic level at which correlation test was performed, reporting correlation (r) and p-value (p)

### Age associations with beta diversity are concordant across taxonomic classifiers

We calculated beta diversity with Bray-Curtis dissimilarity index for each cohort and both taxonomic classifiers to examine the changes in beta diversity with age. We then performed principal coordinate analysis (PCoA) to visualize the similarities and differences between samples (**Figure 3, Figure S5**) (**Table S3-S4**). The percent of variability explained by the first component of the PCoA was greater than 69% at the phylum level and decreased to less than 20% at the species level across classifiers and cohorts. The PCoA plots display greater similarity between samples of similar ages compared to samples of different ages, with statistically significant associations observed in most cases between different classifiers and cohorts (**Figure 3A-B, Figure S5A-B**). One exception included differences at the phylum level in the ILO cohort, in which MetaPhlAn4 did not identify statistically significant changes in beta diversity with age (PERMANOVA F=1.63, p=0.188), although statistically significant differences were observed when using Bracken (PERMANOVA F=4.63, p=0.013) (**Figure 3A**).

**Figure 3.**
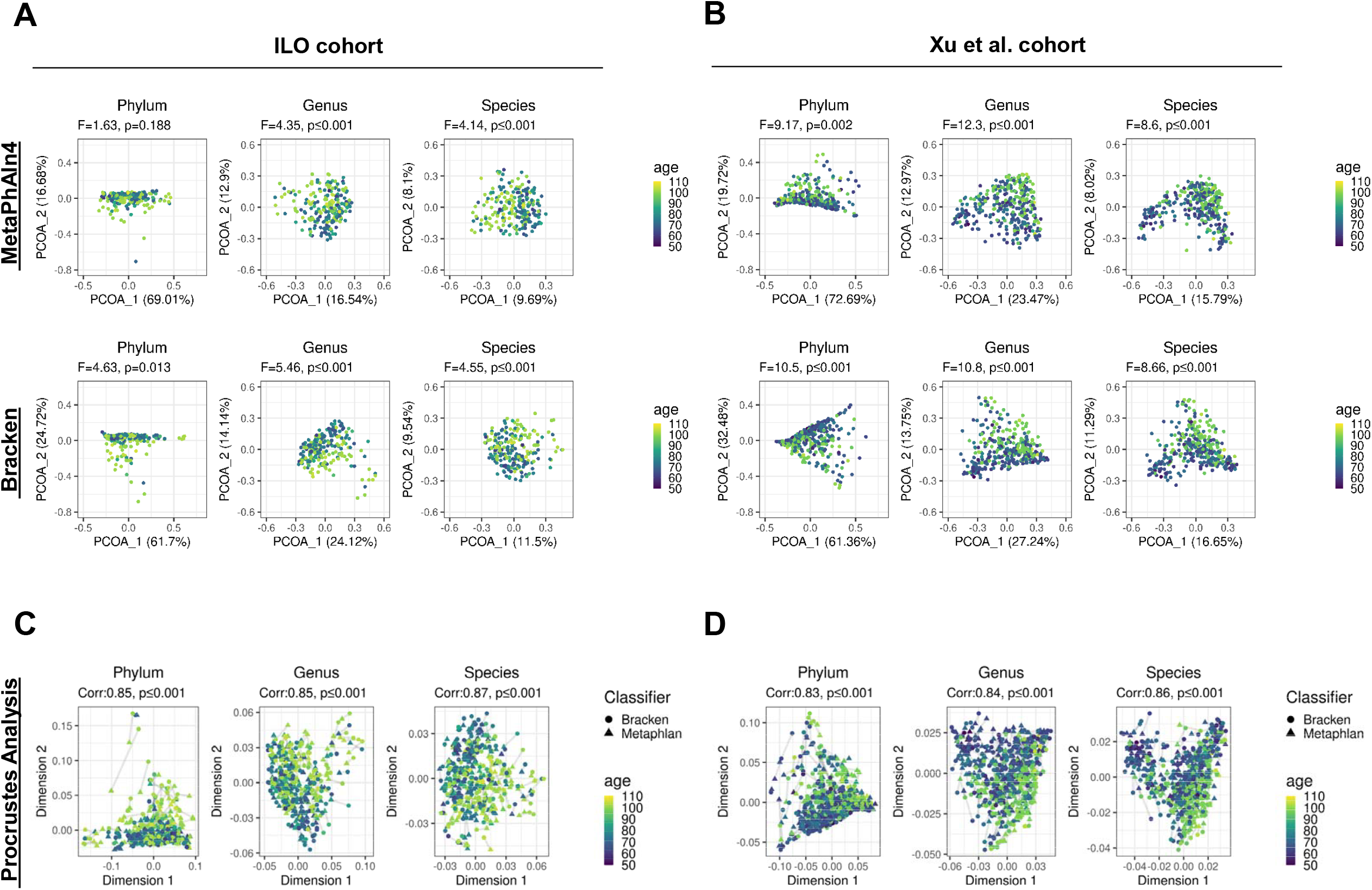
Beta diversity displays similar changes with age based on profiler approach in each cohort. **(A-B)** Principal coordinate analysis plots displaying the Bray-Curtis dissimilarities between samples, comparing across phylum, genus, and species taxonomic levels within cohorts and between cohorts and classification methods. We employed PERMANOVA to evaluate the differences in the Bray-Curtis dissimilarities between samples with the association with age, with a significance threshold set at 0.05. **(C-D)** Scores from Procrustes analysis performed on Bray-Curtis dissimilarities of samples from both classification methods within each cohort, with lines connecting the same samples. We employed Procrustian randomization (Monte Carlo) test to evaluate whether the concordance between the distances based on the taxonomic classifiers is greater than expected due to chance, with significance threshold set at 0.05.

We used Procrustes analysis to show graphically whether the dissimilarity between any pair of samples within a cohort are maintained regardless of the classifier used. To assess this further, we used a Procrustean randomization test to measure whether the correspondence across methods was greater than would be expected due to chance (**Figure 3C-D, Figure S5C-D**). There was high correspondence in dissimilarities at all taxonomic levels (correlation: 0.83-0.87, p≤0.001) between Bracken-based profiles versus MetaPhlAn4-based profiles in both cohorts. Additionally, Mantel tests showed high correlation among Bray-Curtis dissimilarities based on MetaPhlAn4 versus Kraken2 (correlation>0.75, p≤0.001 for all taxonomic levels; **Table 3**). These findings were similar when limiting the analysis to in-common taxa identified by both classifier methods (**Figure S6**). Overall, the beta diversity results suggest that both taxonomic classifiers capture similar community-level changes with age in both cohorts.

### Differential abundance analysis reveals benefits of a multi-classifier approach

To explore changes in individual taxa abundance with age based on each taxonomic classifier, we performed differential abundance analysis at the species level (**Figure 4, Table S5**). The volcano plots in **Figure 4** summarize the species-level results for ILO (**Figure 4A**) and Xu et al. cohorts (**Figure 4B**). In ILO, the analysis based on MetaPhlAn4 identified 55 species whose relative abundances were associated with age, while the analysis based on Kraken2/Bracken identified 59 species associated with age at 5% FDR (**Figure 4C**). In Xu et al., the analysis based on MetaPhlAn4 identified 49 species whose relative abundance were associated with age, while the analysis based on Kraken2/Bracken identified 29 species associated with age at 5% FDR (**Figure 4D**). Use of both MetaPhlAn4 and Bracken resulted in the identification of many age-associated species that would not be found using one method alone. For instance, 43 (21) species in ILO (Xu) were age-associated in Bracken alone, which would not have been identified using MetaPhlAn4, and likewise 39 (41) species in ILO (Xu) were age-associated in MetaPhlAn4 alone (**Figure 4C-D**).

**Figure 4.**
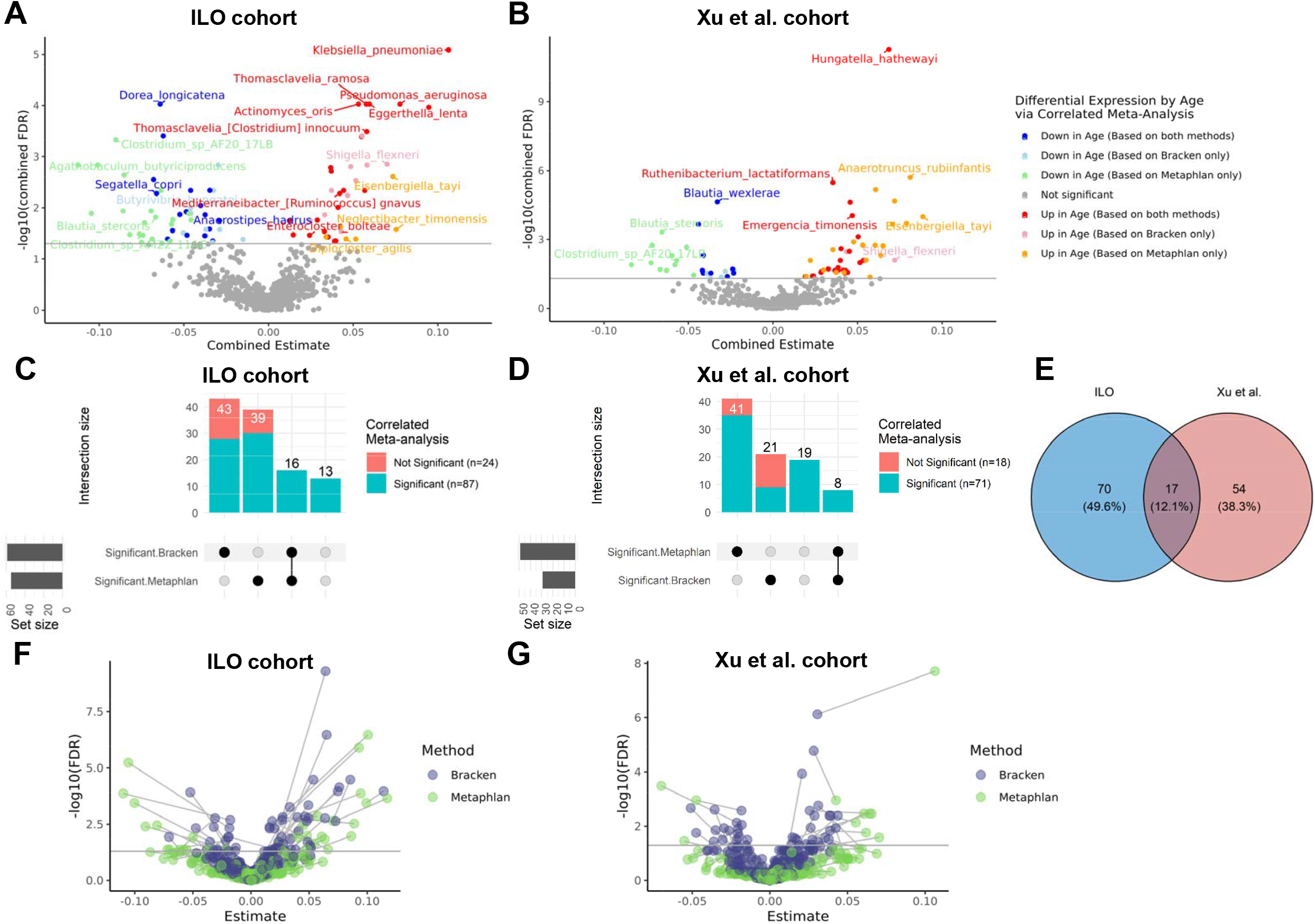
Differential abundance analysis displays shared, classifier-specific and cohort-specific differences. **(A-B)** Volcano plot of the differentially abundant species with age in each cohort. We employed linear regression models with generalized estimating equations to assess age associations with species relative abundance, and used correlated meta-analysis to generate combined p-values and combined effect estimates for species identified by both classifiers within a cohort, with significance assessed with combined FDR < 0.05. Significant differentially abundant species labelled by upregulation and downregulation with age using both taxonomic classifiers or a single classifier. **(C-D)** UpSet plots containing all species significantly associated with age via individual tests or correlated meta-analysis approach. The vertical bar heights show the number of species associated in the Bracken profiles, MetaPhlAn4 (MetaPhlAn) profiles, both, or neither based on individual tests. Blue shading shows the number of species significantly associated with age via the correlated meta-analysis approach, and red shading shows number of species significantly associated with age via individual tests that were not significant using the correlated meta-analysis approach. Of the 87 (71) species identified by the correlated meta-analysis approach in the ILO (Xu) cohorts, only 16 (8) would have been discovered by a single-profiler approach regardless of whether Bracken or MetaPhlAn4 was used. Additionally, 13 (19) age-associated species were identified in the ILO (Xu) cohorts by correlated meta-analysis that would not be identified by either Bracken or MetaPhlAn4 alone. **(E)** Venn diagram of age-associated species identified by the correlated meta-analysis approach in the ILO and Xu et al. cohorts. **(F-G)** Volcano plots of the differentially abundant species with age in each cohort based on individual tests. We employed linear regression models with generalized estimating equations to assess age associations with species relative abundance adjusting for sex and education using both classifiers within a cohort. For each taxon, the estimates and FDR values based on each classifier method are displayed with a line connecting between methods: Bracken (purple) and MetaPhlAn4 (Metaphlan) (green). For each classifier method, significance was assessed with FDR < 0.05.

We used a novel correlated meta-analysis approach^27^ to integrate results across both classifier methods while accounting for the non-independence of tests of association for taxa measured via two classifiers in the same individuals. This enabled the identification of species that were robustly associated with age (i.e. age-associated in both cohorts), including some that would not have been found to be age-associated by a single profiling method. Of the 43(21) species in ILO(Xu) that were significantly age-associated in the Bracken profiles but were not age-associated (or were absent) in the MetaPhlAn4 profiles, 28(9) species displayed significant association with age using the correlated meta-analysis approach (**Figure 4C-D**). Species include *Mediterraneibacter [Ruminococcus] gnavus*, linked to inflammation and aging^28,29^, whose abundance was associated with older age in ILO and Xu et al. cohorts. Of the 39(41) species in ILO(Xu) that were significantly age-associated in the MetaPhlAn4 profiles but were not age-associated (or were absent) in the Bracken profiles, 30 (35) species showed significant association with age by correlated meta-analysis (**Figure 4C-D**). This set of species include *Neglectibacter timonensis, Eisenbergiella tayi*, and *Clostridium_sp_AF20_17LB* that have been previously reported in association with longevity.^30^ In addition, 16 (8) species in ILO (Xu) were found to be significantly associated with age by both classifier methods via individual tests, all of which were also identified by the correlated meta-analysis method (**Figure 4C-D**). Further, correlated meta-analysis also identified species that were not significantly associated with age in *either* MetaPhlAn4 or Bracken individually, but were found to be significantly age-associated in the meta-analysis (13 in ILO, 19 in Xu et al.; **Figure 4C-D**). These include species with documented age associations such as *Anaerostipes hadrus, Faecalibacterium prausnitzii*, and *Akkermansia muciniphila*.^31-35^

Of the 141 total species found to be age-associated in at least one cohort by the correlated meta-analysis, we found 17 were associated with age in both cohorts (**Figure 4E, Table S5**). Finally, we further compared species-level effects and FDR q-values between the classifier methods based on individual tests (**Figure 4F-G**). Age effect sizes tended to be larger for MetaPhlAn4 compared with Bracken, and no clear pattern was present with respect to relative FDR q-values between the classifiers.

## DISCUSSION

### Overview

Previous benchmarking studies of taxonomic classifiers of metagenomic data have advocated for consensus methods as a way of identifying results.^14,17,18^ In this report, we present an analysis of age-associated features of gut microbial communities in older and long-lived adults (50-107 years), in which we use consensus and a novel meta-analytic approach to integrate findings from two popular metagenomic taxonomic classifiers, MetaPhlAn4 and Kraken2. We performed diversity analyses in which both classifiers captured similar age-associated changes in two long-lived cohorts, with species-level variability observed across classifiers and cohorts. We also conducted a novel analysis to integrate differential abundance results between classifiers using a correlated meta-analysis approach, which identified 17 species with robust association with age across cohorts, including species that would not have been detected using one profiler alone.

### Database differences and low-abundance taxa drive profile differences between classifiers

The general observation from examining taxonomic overlap is that the choice of the taxonomic classifier affects in a substantial way the taxa identified at all levels. In terms of species identified in the two cohort studies, Kraken2/Bracken identified 33-67% more species than MetaPhlAn4 did (**Table 1**), which matches findings in previous benchmarking studies.^16^ This difference did not appear to be the result of differences in numbers of species present in the databases, as Kraken2’s database had just 11% more species than MetaPhlAn4’s, and because Kraken2 identified many species that were present in the MetaPhlAn4 database but not identified by MetaPhlAn4 (**Table 2**). This finding also agrees with previous work that characterized Kraken2/Bracken as being more sensitive and more prone to false positive calls than MetaPhlAn4.^12,15^ There are 8 overlapping species among the top 10 abundant species measured by the two classifiers (**Figure S2C-D**), suggesting that there is general agreement as to the most abundant species in the samples from the two methods.

A notable observation is that the reference databases available for MetaPhlAn4 and Kraken2 have relatively few (∼30%) species in-common even when we compare domains that both tools are designed to detect (i.e., excluding viruses). The difference of the reference databases implies that there are thousands of species that can only be identified by one of the tools. These differences may be limitations of the approaches used by each taxonomic profiler. For instance, some species may not have appropriate marker genes to distinguish them from other species using MetaPhlAn4, and therefore may not be included in the curated database for this tool. On the other hand, the MetaPhlAn4 database includes many species genome bins (SGBs) from metagenome-assembled genomes, expanding on the offerings available compared to the Kraken2 RefSeq databases.

### Age trends in microbiome diversity generally display correspondence across classifier methods

Despite the different microbiome compositions detected by the two tools, the downstream microbiome diversity analyses generally pointed to similar findings. For example, the associations of the normalized alpha diversity with age were comparable between classifiers at most taxonomic levels, with some disagreement at the species level (**Figure 2**). Both classifiers identified that alpha diversity in the gut microbiome increased with age at the phylum level, a finding which agrees with previous work.^23,36,37^ In addition, we observed strong correlations between the normalized alpha diversity scores calculated from both classifiers’ profiles, suggesting the samples generated with different taxonomic classifiers have comparable distributions of taxa in their taxonomic profiles. Despite these strong correlations, inconsistencies in species-level alpha diversity associations in particular highlight the existence of classifier-specific biases not only in which species are identified (**Figure 2A-B**), but in quantifying relative abundances of the detected species, with downstream implications for biological interpretations. Other studies have also found inconsistencies in age-related changes in species-level alpha diversity^24^, and our observations suggest that choice of taxonomic classifier may be a factor contributing to such inconsistencies.

Beta diversity ordination plots revealed similar relationships among samples when using MetaPhlAn4 versus Kraken2/Bracken, and comparable associations between taxonomic composition and age based on PERMANOVA (**Figure 3**). The significant changes in beta diversity with age observed using both classifiers have also been documented in previous studies. ^23,24,39,40^ Procrustes analysis and Mantel tests showed significant correspondence and high correlation of the taxonomic profiles at all levels (**Figure 3C-D, Table 3**), suggesting that despite taxon-level differences in profiles, broad phenotypic associations with taxonomic profiles may be preserved across classifiers.

### Differential abundance analyses using both classifier methods capture more age-associated taxa

Consensus and meta-analytic approaches may be especially fruitful when comparing results from phenotype associations with individual taxa (**Figure 4, Table S5**). Individual taxa are subject to false positive identifications, false negatives, and biases in quantification, and thus tools to identify robust associations are critical.^16^ To synthesize evidence from across classifier methods, we applied a correlated meta-analytic procedure currently in development by our group^27^ to combine p-values from age association tests performed on two taxonomic profiles from the same samples. In cases where only one method identified a given taxon, the p-value from the individual test was used. Of the 141 species found to be associated with age using this method, 17 were significantly age associated in both cohorts, representing the taxa with the greatest evidence for age association based on our data.

Among these 17 age-associated taxa with replication across cohorts, some taxa would have been missed if we used only one taxonomic classifier. Among taxa that were only identified by MetaPhlAn4, *Neglectibacter timonensis, Eisenbergielle tayi*, and *Clostridium_sp_AF20_17LB* were identified as consistently different in long-lived individuals compared to younger individuals across cohorts in a study that included the Xu et al. cohort and seven other cohorts.^30^ For other species identified by MetaPhlAn4 alone, including *Diplocloster agilis, Clostridium sp_AM22_11AC, Blautia stercoris, Agathobaculum butyriciproducens*, we did not find previously documented associations with aging or longevity. However, *A. butyriciproducens* and *B. stercoris*, which we found to be age-associated in both cohorts in our study, have been linked to mouse models of age-related changes in cognition and other conditions. *A. butyriciproducens* was found to decrease age-associated cognitive deficits ^41^ and AD-related cognitive deficits/pathology^42^ in mice. *A. butyriciproducens* was found to have neuroprotective effects in mouse models of Parkinsons,^43^ and has been associated with decreased PET amyloid burden in humans with Alzheimer’s disease.^44^ *B. stercoris* has also been used in mouse models of autism spectrum disorder to decrease behavioral deficits.^45^

Among taxa that were only identified by Kraken2/Bracken, *Mediterraneibacter [Ruminococcus] gnavus* was found to be significantly associated with age in both cohorts in our study. *M. gnavus* has been previously linked to gut dysbiosis and inflammatory conditions such as irritable bowel syndrome, relevant to aging.^28,46^ In addition, *Shigella flexneri* was found to have significant age-association in both cohorts in our study when using only Kraken2/Bracken. This finding, based on a single classifier approach, may be explained by the fact that *S. flexneri* is very closely related to *Escherichia coli* and the species can be hard to differentiate, with previous work showing that k-mer based methods may be effective at differentiating *S. flexneri* while marker-gene based methods such as MetaPhlAn4 are unable to do so.^47,48^ Other age-associated taxa identified by Kraken2/Bracken alone were very low abundance species with no previous isolates in humans (*Butyrivibrio hungatei*, generally found in ruminant animals,^49^ and *Romboutsia ilealis*, isolated from the small intestine of the rat^50^). It is possible Kraken2/Bracken is mis-identifying reads from human-associated close relatives as originating from these species, given the known tendency of the k-mer based tools to generate false positive calls, especially for low abundance species. While considering the union of age-associated species across profilers allows for a greater number of age associated species to be identified, it is important to consider the limitations of each classifier included, and we consider species identified by both classifiers to reflect the most likely true positive associations.

We also identified species that were significantly associated with age by the correlated meta-analysis, but not by individual tests. These were generally taxa with borderline significance in the individual tests from both profiles (i.e. p-values near but greater than 0.05), where the concordance in associations across profiles resulted in a significant meta-analyzed p-value. These associations include species previously associated with inflammation and aging (e.g. *Akkermansia muciniphila*^*31*^) and species that produce anti-inflammatory short-chain fatty acids (SCFA) (e.g. *Anaerostipes hadrus, Faecalibacterium prausnitzii*). Inflammation highly impacts aging, as chronic, low-grade inflammation is considered a hallmark of aging.^51^ The genus *Akkermansia* has been shown to be highly abundant in long-lived individuals,^52^ and a study of *A. muciniphila* in mouse models showed the decline of age-related changes in older mice.^32,33^ *A. hadrus* and *F. prautsnitzii* are considered beneficial bacteria and producers of butyrate which have anti-inflammatory properties. A decrease in butyrate-producing bacteria has been shown to increase inflammation and is associated with inflammatory diseases such as inflammatory bowel disease.^34,35^ In addition, strains of *Phocaeicola vulgatus*, another species identified as age-associated by meta-analysis in our study, have also been shown to affect inflammatory diseases.^53^ *P. vulgatus* is a highly abundant bacterium in the human gut that degrades complex carbohydrates to produce SFCAs, reducing inflammation. In summary, the meta-analytic approach we employed to combine results between the taxonomic classifiers allowed us to identify taxa with biologically plausible age associations that would not have been otherwise identified.

### Limitations of the study

A limitation of this study is that, when comparing taxonomic profilers on real-world data, there is no “ground truth” or gold standard available for comparison. We focus instead on agreement and disagreement between the methods, as well as robustness of associations across cohorts and, to a degree, previous literature support for age associations. Another limitation is that our data is cross-sectional, such that we cannot make inferences about aging and longevity processes within individuals. Additionally, we restricted our analysis to taxa mappable to the NCBI database to compare and integrate across profilers, which excludes a large number of SGB-based taxonomic assignments available in MetaPhlAn4 in particular. However, the MetaPhlAn4 paper also used this approach to benchmark MetaPhlAn4 against other classifiers.^2^ Furthermore, for brevity, we used one type of alpha diversity and beta diversity measure to compare age association changes in diversity of microbial communities across classifiers and cohorts. We intentionally used the normalized alpha diversity metric to assess age association changes across taxonomic levels as well as across classifier methods and cohorts. In addition, we selected Bray-Curtis dissimilarity index to measure beta diversity as it is a highly utilized measure in microbiome studies of aging and longevity. Finally, a limitation of the correlated meta-analysis approach is that it may identify taxa as significant although they may be insignificant by one method and missing by another method. However, this was observed for only 2 species in our study that became borderline significant via correlated meta-analysis approach due to minimal differences introduced by the different FDR calculations within methods.

### Conclusions

In this study, we highlight the utility of integrating results from multiple classification methods when performing downstream analysis of microbiome data with phenotypes of interest, by analyzing data from a new Integrative Longevity Omics (ILO) cohort and a replication cohort. Previous work has highlighted the respective strengths of various taxonomic classification methods. Using two popular classifiers, we show that analyses such as Procrustes and a novel correlated meta-analytic procedure can highlight reproducible associations with biological plausibility. We also show that, regardless of the method chosen, using a single taxonomic classifier (as is standard practice in the field) results in missing potentially meaningful taxonomic associations with phenotypes.

## RESOURCE AVAILABILITY

### Lead contact

Further information and requests for resources and reagents should be directed to and will be fulfilled by the lead contact, Tanya Karagiannis

### Materials availability

This study did not generate new unique reagents, and all materials used in this study are reported either in the main text or in the supplemental information.

### Data and code availability

- Shotgun metagenomics sequencing data from Xu et al. cohort^25^ is publicly available and was accessed from the NCBI BioProject repository (acquisition number: PRJNA613947)
- Shotgun metagenomics sequencing data from the ILO cohort will be available on the ELITE Portal: https://eliteportal.synapse.org
- The pipeline scripts for metagenomics sequencing processing are available on github: https://github.com/Integrative-Longevity-Omics/MGS_pipeline
- All original analysis scripts are available on github: https://github.com/Integrative-Longevity-Omics/IntegrativeTaxonomicAnalysis

## ACKNOWLEDGMENTS

TK, YC, SB, AT, SM, DS, SA, TP, PS, MS are supported by NIH-NIA UH2/UH3AG064704. NIH S10OD032203 shared instrumentation grant for AT at Tufts University Core Facility Genomics Core.

## AUTHOR CONTRIBUTIONS

TK, DS, PS and MS conceived the analyses. TP, PS established the ILO study and procured funding. AT carried out experiments. TK, YC, SB, and MS analyzed the data. TK and MS wrote the paper, and all authors provided feedback on the manuscript.

## SUPPLEMENTAL INFORMATION

**Document S1. Figure S1-S6**

**Document S2. Supplementary Materials and Methods**.

**Table S1. Number of taxa identified by MetaPhlAn4 and Kraken2/Bracken across taxonomic levels**.

**Table S2. Normalized alpha diversity scores for samples in each cohort based on MetaPhlAn4 and Kraken2/Bracken profiles across taxonomic levels**.

**Table S3. Beta diversity PCoA components for samples in ILO cohort based on MetaPhlAn4 and Kraken2/Bracken profiles across taxonomic levels**.

**Table S4. Beta diversity PCoA components for samples in Xu cohort based on MetaPhlAn4 and Kraken2/Bracken profiles across taxonomic levels**.

**Table S5. Differential abundance analysis results based on individual tests and correlated meta-analysis approach for each cohort**.

