## Supplementary material for "Integrative analysis across metagenomic taxonomic classifiers: A case study of the gut microbiome in aging and longevity": Figure S1-S6

### Slide 1
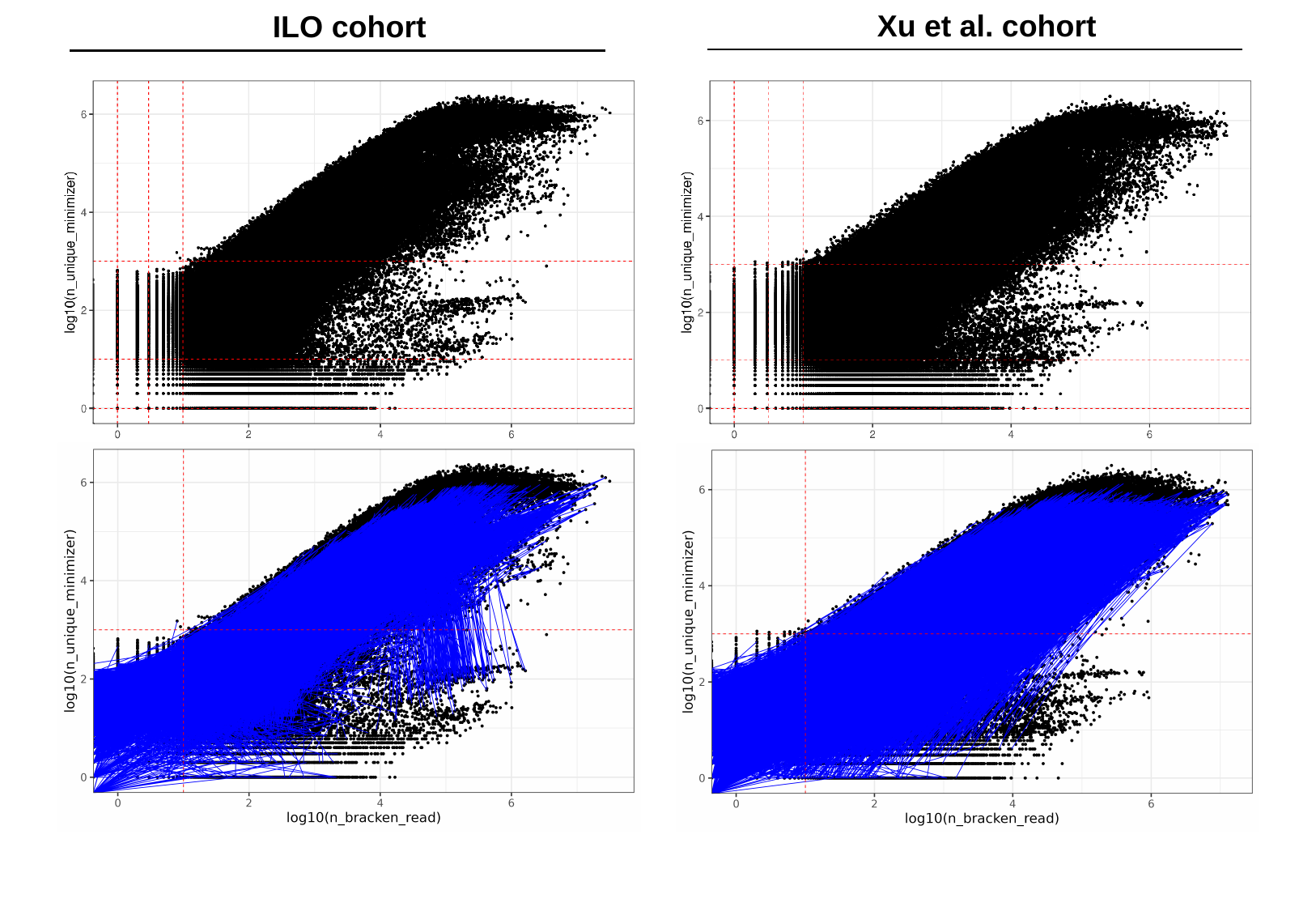

Xu et al. cohort
 ILO cohort

### Slide 2
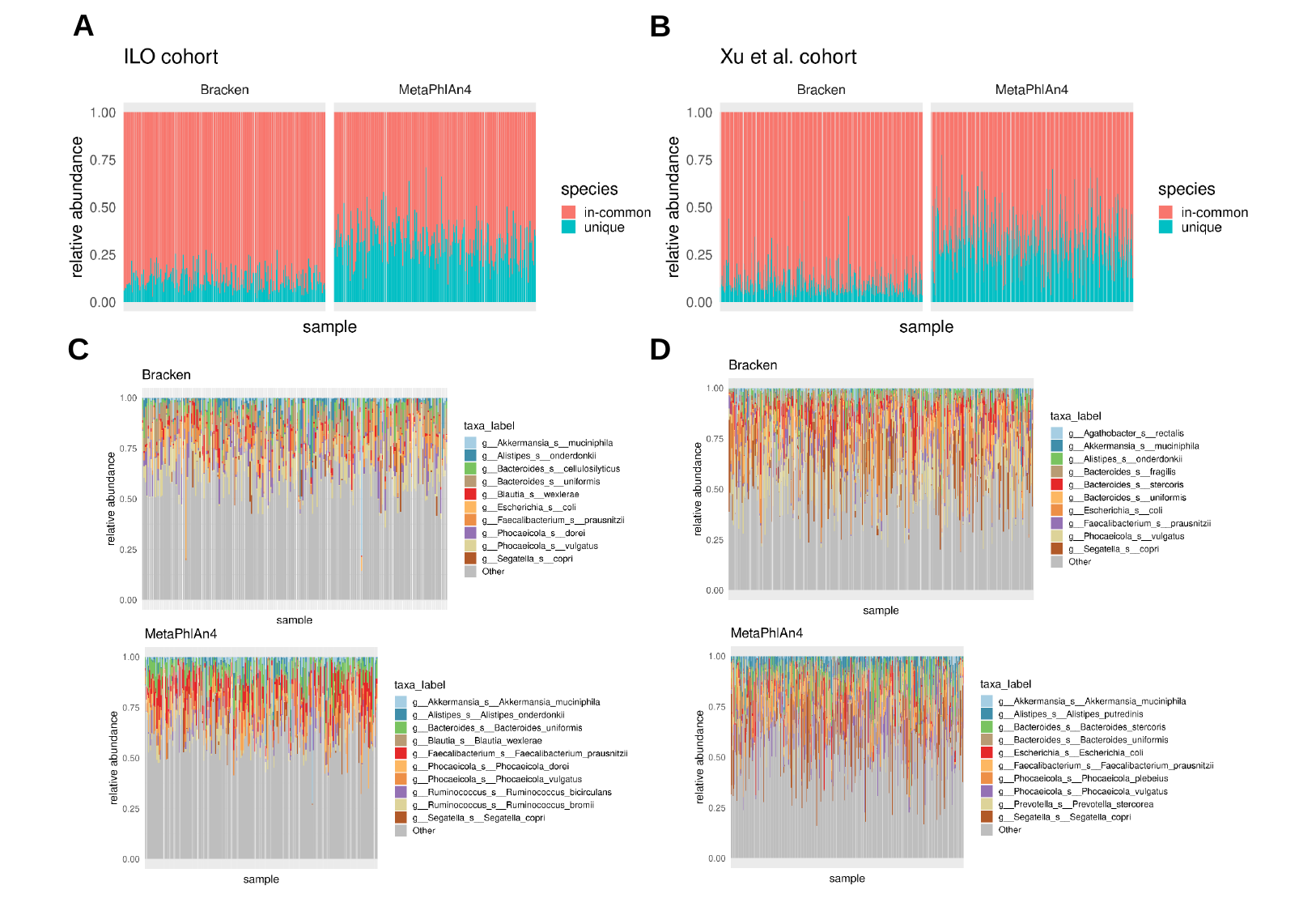

A
B
D
C

### Slide 3
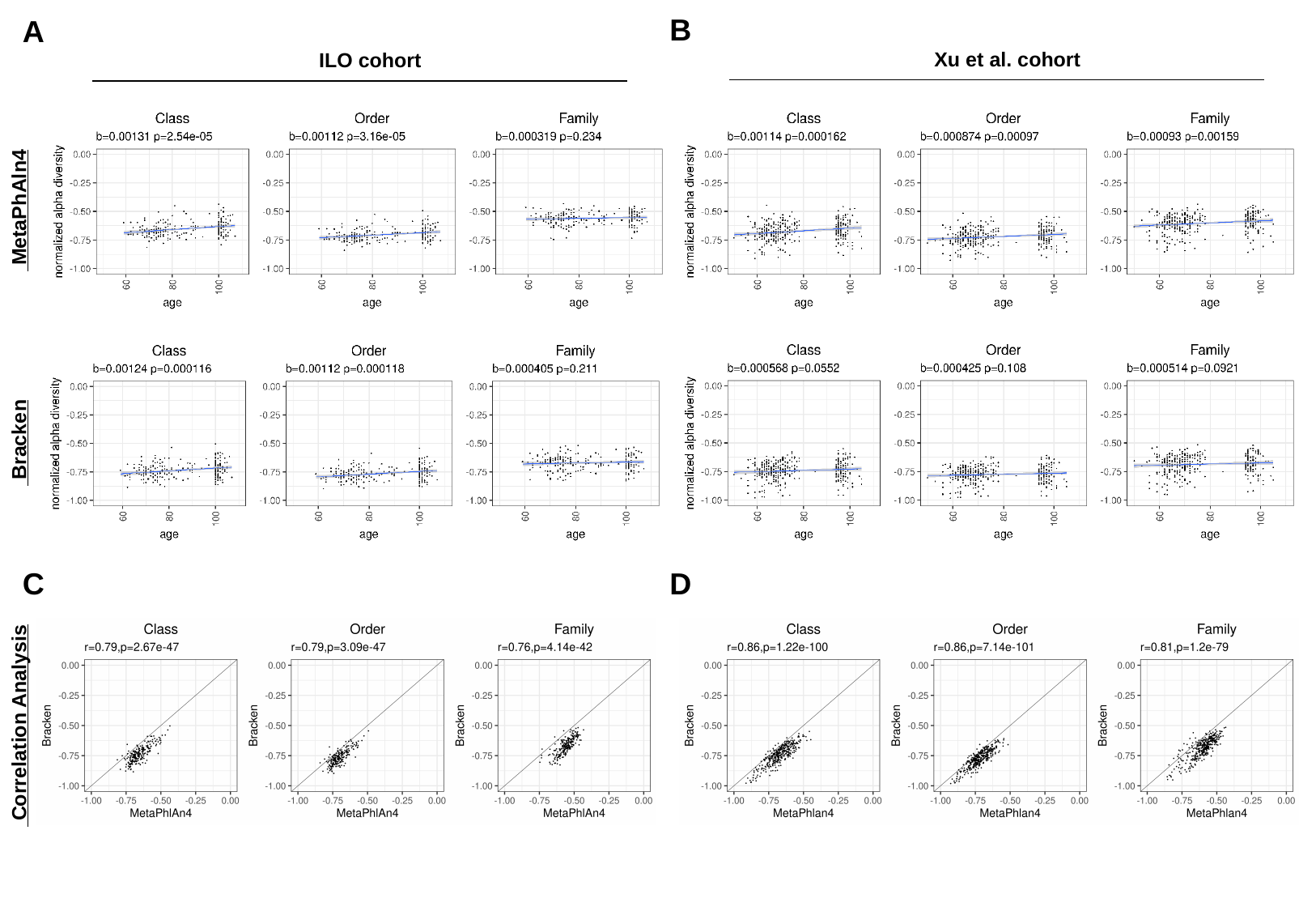

B
A
 Xu et al. cohort
 ILO cohort
 MetaPhAln4
 Bracken
C
D
 Correlation Analysis

### Slide 4
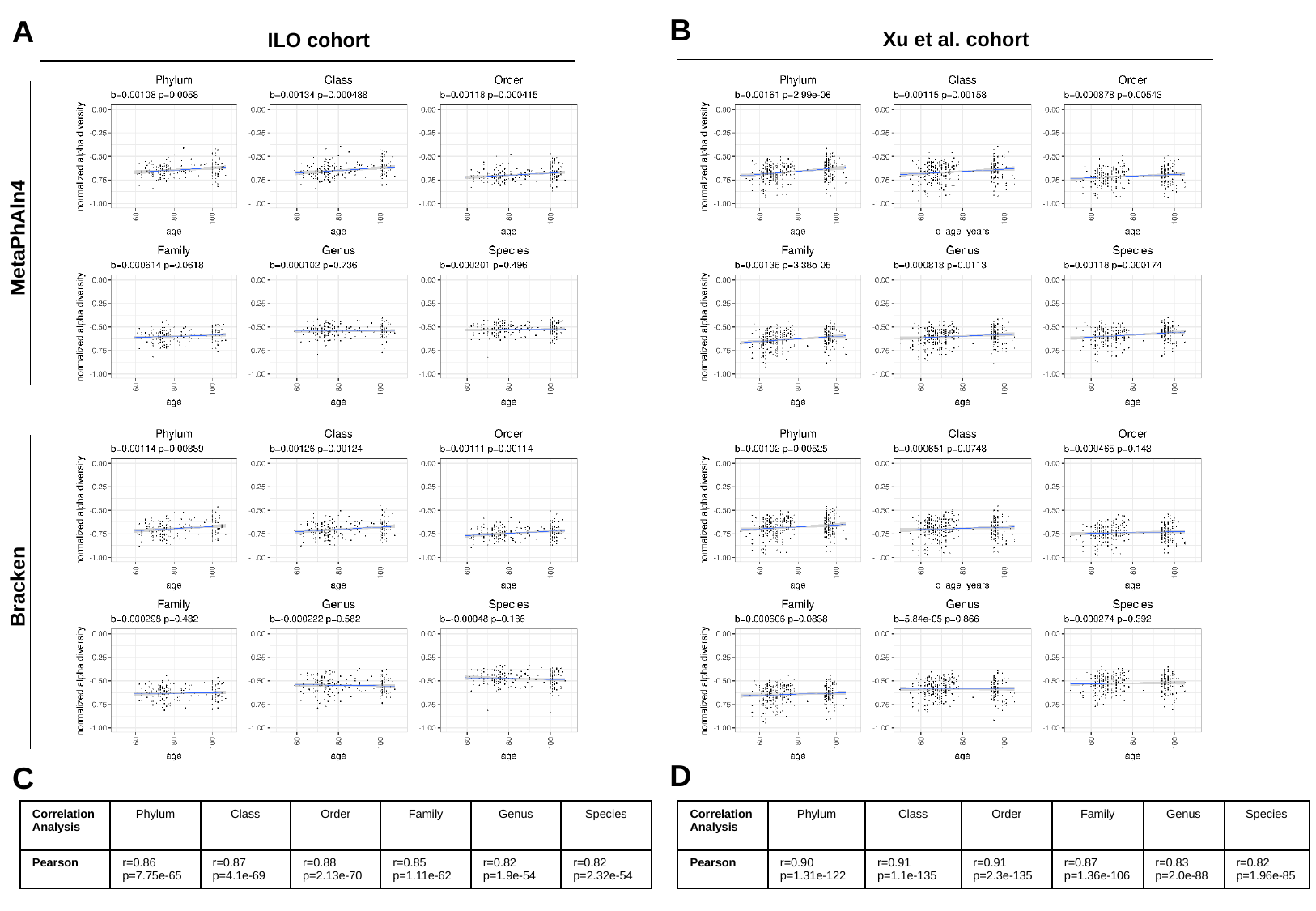

B
A
 Xu et al. cohort
 ILO cohort
 MetaPhAln4
 Bracken
D
C
| Correlation Analysis | Phylum | Class | Order | Family | Genus | Species |
| --- | --- | --- | --- | --- | --- | --- |
| Pearson | r=0.86 p=7.75e-65 | r=0.87 p=4.1e-69 | r=0.88 p=2.13e-70 | r=0.85 p=1.11e-62 | r=0.82 p=1.9e-54 | r=0.82 p=2.32e-54 |
| Correlation Analysis | Phylum | Class | Order | Family | Genus | Species |
| --- | --- | --- | --- | --- | --- | --- |
| Pearson | r=0.90 p=1.31e-122 | r=0.91 p=1.1e-135 | r=0.91 p=2.3e-135 | r=0.87 p=1.36e-106 | r=0.83 p=2.0e-88 | r=0.82 p=1.96e-85 |

### Slide 5
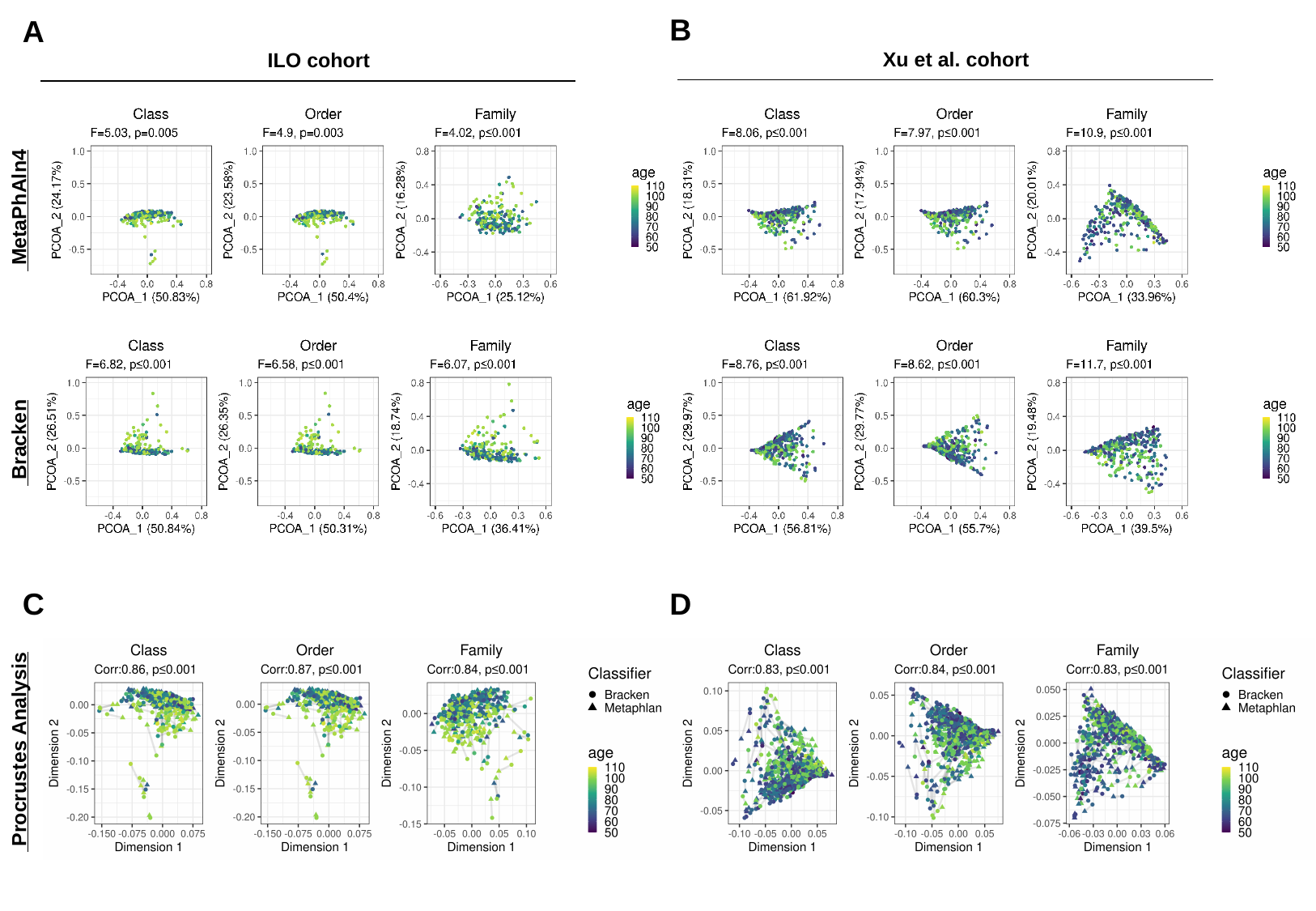

B
A
 Xu et al. cohort
 ILO cohort
 MetaPhAln4
 Bracken
C
D
 Procrustes Analysis

### Slide 6
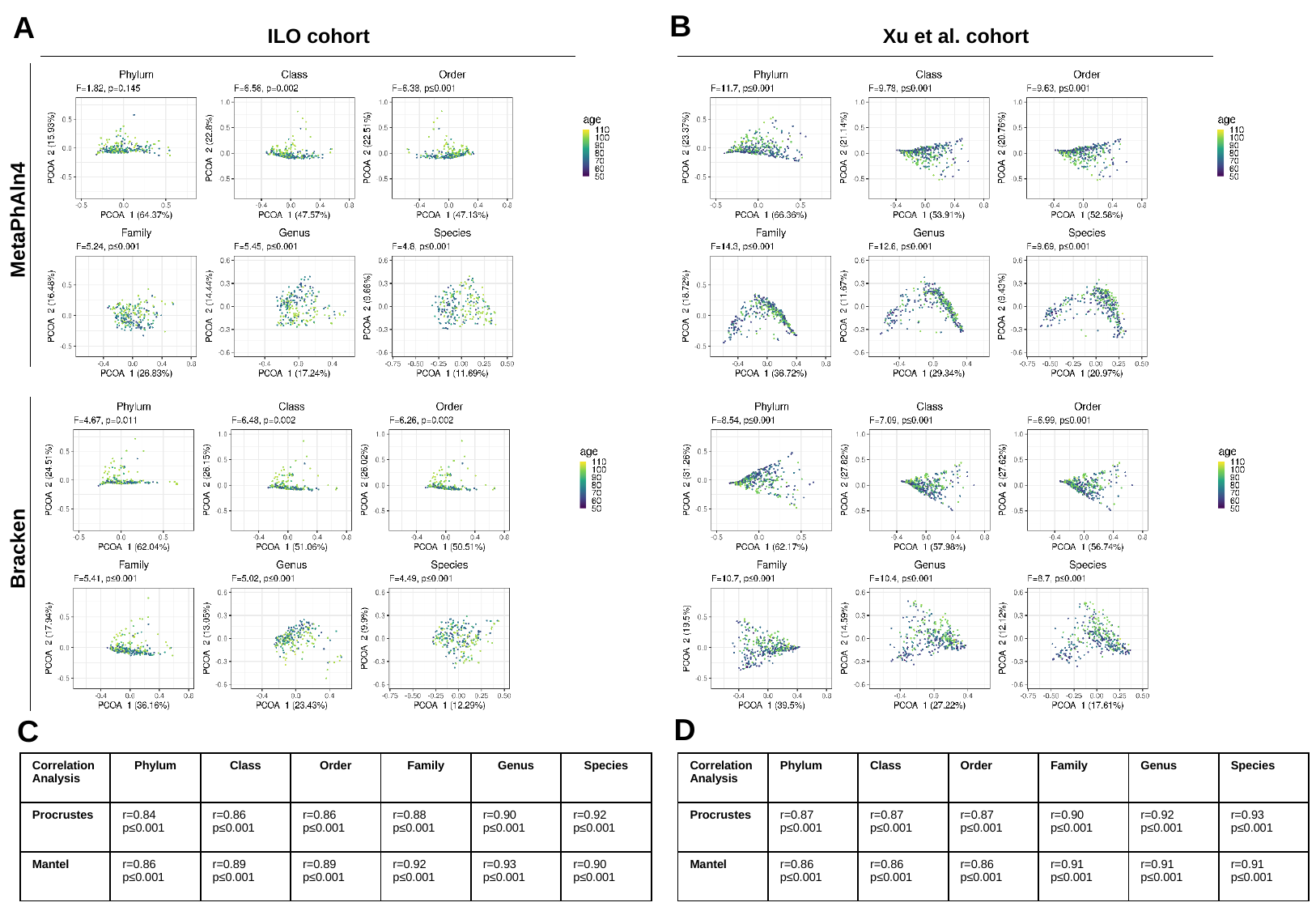

B
A
 ILO cohort
 Xu et al. cohort
 MetaPhAln4
 Bracken
D
C
| Correlation Analysis | Phylum | Class | Order | Family | Genus | Species |
| --- | --- | --- | --- | --- | --- | --- |
| Procrustes | r=0.84 p≤0.001 | r=0.86 p≤0.001 | r=0.86 p≤0.001 | r=0.88 p≤0.001 | r=0.90 p≤0.001 | r=0.92 p≤0.001 |
| Mantel | r=0.86 p≤0.001 | r=0.89 p≤0.001 | r=0.89 p≤0.001 | r=0.92 p≤0.001 | r=0.93 p≤0.001 | r=0.90 p≤0.001 |
| Correlation Analysis | Phylum | Class | Order | Family | Genus | Species |
| --- | --- | --- | --- | --- | --- | --- |
| Procrustes | r=0.87 p≤0.001 | r=0.87 p≤0.001 | r=0.87 p≤0.001 | r=0.90 p≤0.001 | r=0.92 p≤0.001 | r=0.93 p≤0.001 |
| Mantel | r=0.86 p≤0.001 | r=0.86 p≤0.001 | r=0.86 p≤0.001 | r=0.91 p≤0.001 | r=0.91 p≤0.001 | r=0.91 p≤0.001 |
