## Supplementary Materials and Methods for "Integrative analysis across metagenomic taxonomic classifiers: A case study of the gut microbiome in aging and longevity"

### Supplementary Methods

#### Experimental Procedure:

<u>Recruitment of human subjects.</u>

Integrative Longevity Omics (ILO) study: The ILO study enrolled ~1400 centenarians, their biological offspring, and spouses of the offspring (a referent cohort) between 2019 and 2024 from North America. Study participants complete a comprehensive assessment of physical and cognitive function, answer questionnaires on medical history and medication use, and provide blood and stool samples for genetics and genomic studies. Data collection uses a combination of phone interviews, questionnaires, virtual and in-home visits. Centenarians provided documentation of date of birth. All participants provided written, informed consent and the study and procedures are approved and overseen by a single Institutional Review Board at Albert Einstein College of Medicine. We included the first 220 ILO participants whose stool samples were received, comprising 149 females and 70 males with an average age of 85 years at enrollment, and range 59 to 107 years. All participants identified as being of North American/European descent. Among these participants, 2 identified as African American, 1 identified as Asian American, and the others as White. Age at stool sample collection was calculated based on date of stool sample collection and date of birth. All downstream analyses use age at stool sample collection to represent age. All phenotypic data was collected in REDCap and we used the data export from November 2024.

<u>Stool sample collection.</u>

Participants provide stool samples that they collect using the OMNIgene GUT sample collection kit. Study participants complete a questionnaire accompanying the stool sample collection that includes specific food/supplements consumed in the a) 24 hours and b) one week preceding the sampling. Participants use the self-collection kit to obtain the stool sample and ship it to the genetics core at Boston University for storage. Samples were frozen at –80ºC, except for the first phase of 36 samples which were stored at room temperature.

<u>Shotgun metagenomics sequencing of the gut microbiome of ILO subjects.</u>

DNA from the stool samples was extracted at the Division of Geographic Medicine and Infectious Diseases. Library preparation and next generation sequencing were performed in the Tufts University Core Facility Genomics Core. From each sample, 100 – 500ng of DNA was used as input for preparation of sequencing library using Illumina Nextera DNA Prep kit or Illumina DNA Prep kit. Sequencing for the first phase of samples (36 samples) was performed using Illumina HiSeq platform with High Output V4 chemistry and150-bp paired-end read length. Sequencing for the second phase of samples (184 samples) was performed using Illumina NovaSeq 6000 platform with S4 300 cycle v1.5 chemistry and 150-bp paired-end read length to 10Gbases/sample targeted depth.

Xu et al. replication cohort.

We downloaded the shotgun metagenomics sequencing data of the gut microbiome of 348 individuals from Qidong, Jiangsu Province, China, as paired-end FASTQ files from the NCBI BioProject repository (acquisition number: PRJNA613947)[1]. Clinical metadata from participants was also available. Sequencing was performed using the Illumina HiSeq 4000 platform with a 150-bp paired-end read length to a mean depth of 7.6 Gb. The data set included 178 females and 170 males with an average age of 77 years at enrollment, and range 50 to 105 years. All participants identified as being of Han Chinese descent.

**Unified Preprocessing and Taxonomic Classification:**

We applied the same processing and taxonomic classification as described below for the shotgun metagenomics sequencing data in ILO cohort and Xu et al. cohorts. Sequence data was run through the KneadData software from the Biobakery suite of tools for metagenomic analysis, and the resulting quality-controlled reads were classified using 1) MetaPhlan4 for marker-gene-based taxonomic/functional profiling, and 2) Kraken2 followed by Bracken for kmer-based DNA read classification and subsequent estimation of species abundances from Kraken-classified read profiles using a Bayesian probabilistic model. Details on steps are given below, and the pipeline and documentation are available here: https://github.com/Integrative-Longevity-Omics/MGS_pipeline.

Sequence QC.

We pre-processed FASTQ sequences with KneadData (v.0.12.0) prior to taxonomic classification. We used KneadData to trim adapters and low quality bases using Trimmomatic[2], remove repetitive sequences using Tandem Repeats Finder (TRF)[3], and remove reads from human DNA using bowtie2.[4] We ran KneadData using a bowtie2 database built from the T2T-CHM13v2.0 human reference genome. Unless otherwise specified, everything was run with the default parameters provided by KneadData. We generated sequence quality reports for each of the FASTQ files using FastQC v.0.11.9 (bioinformatics.babraham.ac.uk/projects/fastqc/), and generated a single report using MultiQC v.1.12[5] to inspect the sequence quality of the samples.

Marker-gene-based taxonomic classification.

We used MetaPhlAn4 (v.4.1)[6] for marker-gene-based taxonomic profiling of the shotgun metagenomic sequencing data, with the vJun23 version of its curated marker gene database (https://github.com/biobakery/MetaPhlAn).

K-mer-based taxonomic classification.

We used Kraken2[7] for k-mer-based taxonomic classification. We built a custom Kraken2 database as described by the Kraken protocol[8] to include complete genomes in RefSeq for the bacterial, archaeal, viral, and plasmid domains, along with the human T2T-CHM13v2.0 genome, and collections of eukaryotic pathogens (EuPathDB) and known vectors (UniVec_Core) (downloaded and built September 29, 2023). We used Kraken2 v.2.1.2 with the Kraken2Uniq option to report the k-mer minimizer counts (--report_minimizer_data). We used additional options to set a minimum of 4 overlapping k-mers sharing the same minimizer, and report all taxa including those that have zero read counts. To estimate the taxonomic relative abundances at the species level for each sample, we applied Bracken (v2.9) to the Kraken2 classification report files to compute the estimated species abundances using a Bayesian algorithm.

**Quality Control Pipeline after Taxonomic Classification:**

| QC steps | Description | Kraken2 | Metaphlan4 |
|---|---|---|---|
| Step 1: Initial taxa filtering | Remove taxa with 0 read counts across all samples | X | X |
| | Combined *Prevotella copri* different SGBs with the same NCBI ID into one consensus taxon | | X |
| | Remove taxa that fall under kingdom virus | X | |
| Step 2: Complete classification | Remove taxa with incomplete classifications and keep all taxa classified to species level | X | X |
| | Remove taxa that do not have an NCBI ID assigned | X | X |
| Step 3: Sample QC | Assess samples based on distribution of taxa read counts or relative abundances | X | X |
| Step 4: Taxa QC | Remove likely false positive taxa based on read count and unique k-mer minimizer thresholds | X | |

Initial taxa filtering.

For both classifier datasets, we performed initial filtering to keep only taxa with at least one nonzero abundance value across the samples. For the MetaPhlAn4 processed data, there were no human reads assigned, and only taxa from the kingdom Bacteria, Archaea, and Eukaryota remained. There was only one case where taxa with the same NCBI species identification number were split into separate species-genome-bins (SGBs) within MetaPhlAn4 (*Prevotella copri* is represented by four different SGBs[9]). All SGBs with the same NCBI taxonomic ID were summed together and treated as one to be comparable across profiling methods. For Kraken2/Bracken data, we removed human reads assigned to the NCBI *homo sapiens* species ID 9606. In addition, any taxa under the kingdom Virus within Bracken were removed, with Bacteria, Archaea, and Eukaryota remaining.

Complete classification.

In both the processed MetaPhlAn4 and Kraken2/Bracken dataset. Any taxa that were missing an NCBI identification number for any level of classification at or above the species level were removed from analysis, retaining only taxa with complete NCBI classification.

Sample QC.

We assessed sample quality based on the distribution of taxa read counts and relative abundances. For the Kraken2/Bracken datasets, samples with total read counts deviating more than three standard deviations from the mean were considered outlier samples. For both MetaPhlAn4 and Bracken datasets, samples with a median relative abundance deviating more than three standard deviations from the mean were also considered outlier samples. Furthermore, the distribution of normalized relative abundances across samples was assessed for sample variability due to seasonal changes. Two samples identified as outliers in the

Bracken dataset based on these metrics were selected for further assessment. This step provided insights into the overall composition and variability of the samples. No samples were removed in this analysis to maintain a consistent set of samples across all taxonomic classifiers.

Taxa QC.

Kraken2 has a well-documented tendency to assign false positive taxonomic assignments to reads[10,11], and filtering taxa with low read count or low number of unique k-mer minimizers can help reduce this problem. There are no universal guidelines for choosing threshold values[12], and we examined a range of possible read thresholds (1, 4, and 10) and unique minimizer thresholds (1, 10, and 1,000). A recent pre-print suggests that high correlations between abundances of low-confidence taxa (with few unique minimizers) and high-confidence taxa (greater unique minimizers) can identify potential false positive calls, as the low-confidence taxon is the result of reads from the high-confidence taxon being misclassified.[13] To assess this in our data, we plotted the read counts and unique minimizers present in each taxon within each sample, and connected points corresponding to the greatest correlations among taxa (>0.8; **Supplementary Figure 1**). This revealed low-confidence taxa that were highly correlated with high-confidence taxa, which guided our choice of unique-minimizer threshold value to eliminate as many of these likely false positives as possible. We decided on a read count threshold of 10 and a unique minimizer threshold of 1,000, which also aligns with the Bracken read count default threshold and previously published work.[12]

**Statistical Analysis:**

Association of alpha diversity with age.

To compare changes in alpha diversity with age within each cohort and classification method, we normalized the alpha diversity as:

$$E_s = \frac{-\sum_{i=1}^{k} p_{i_s} log(p_{i_s})}{log(k)} - 1$$

The normalized alpha diversity ($E_s$) is equivalent to the Shannon index, normalized by the number of taxa ($k$) classified at a given taxonomic level in a sample $s$ to make the entropy measure comparable across taxonomic levels as well as across classifier methods and cohorts.[14] The score ranges from -1 and 0, with the minimum score of -1 corresponding to a sample with a single taxon present, and the maximum score of 0 corresponding to a sample with equal proportions of all taxa present in any sample.

We performed linear regression to assess differences in the normalized alpha diversity with ageand used p-value < 0.05 to determine statistical significance. We performed Pearson correlation test to compare sample alpha diversity between the two classification methods within each cohort, and we used a p-value < 0.05 to determine statistical significance.

We applied the normalized alpha diversity as described above to compare the differences in alpha diversity with age when restricting each cohort dataset based on the in-common taxa identified by both methods.

Association of beta diversity with age.

We calculated the Bray-Curtis dissimilarity index between pairs of samples based on the matrix of taxa proportions using the *rbiom* R package (https://github.com/cmmr/rbiom). We then performed Principal coordinate analysis (PCoA) using the *ape* R package (https://github.com/emmanuelparadis/ape) to visualize the similarities and differences between samples based on the Bray-Curtis dissimilarities. We performed PERMANOVA using the *vegan* R package (https://github.com/vegandevs/vegan) with 999 permutations to assess differences in the Bray-Curtis dissimilarities with age, and we used a p-value < 0.05 to determine statistical significance. To compare the differences in beta diversity with age when filtering each cohort dataset based on the in-common taxa identified by both methods, we performed the same steps as described above.

We performed Procrustes analysis to quantify and visualize the concordance in beta diversity between taxonomic classification methods. We used the *procuste* function in the *ade4* R package[15] to generate scaled and rotated coordinates of points from each classifier, and plotted the transformed coordinates with lines connecting points representing the same sample. We used *procuste.rtest* with 999 permutations to calculate and test the correlation between the datasets. Furthermore, we performed Mantel correlation test using the *ape* R package with 999 permutations as an additional statistical test to evaluate the differences in the sample Bray-Curtis dissimilarities between methods.

Although a few significant associations were present between these diversity metrics and experimental phase at particular taxonomic levels, these associations were primarily driven by outlier samples, and no associations were present between experimental phase and covariates of interest including age. Thus, we did not adjust for experimental phase in the statistical analyses in this study.

Differential abundance analyses with age.

We used generalized estimating equations (GEE) with an exchangeable correlation structure from the *geepack* R package[16] to model the relationship between microbial relative abundance and phenotypic variables, while accounting for the potential for within-family clustering. The models included the log-transformed relative abundance values of each taxa as the dependent variable, and age, sex, and education level as covariates. A pseudocount, calculated as half the minimum non-zero value of each taxon across samples, was added to zero values in the original OTU table to allow for log-transformation of these values, similar to what is done in MaAsLin 2 software[17]. The data were then renormalized to relative abundances and log-transformed to enhance normality. Benjamini-Hochberg-adjusted p-values were calculated to control false discovery rates.

Harmonization of differential abundance analyses across taxonomic classifiers.

To harmonize differential abundance analyses incorporating information across classifiers within each cohort, we generated combined p-values and estimates for all species that were identified by both MetaPhlAn 4 and Kraken2. To combine p-values, we used a correlated meta-analysis procedure recently developed in our group to calculate an “Adjusted maximum p-value" for the age association of each species across classifiers[18], which accounts for the non-independence of results being meta-analyzed—in this case, age associations with two measurements of species abundance performed on the same samples. For the combined effect estimates, we used the mean of the effect estimates from the two classifiers, since the sample sizes for classifiers within cohort are equivalent. For species that were identified by one classifier only, we used the p-values and effect estimates from the available classifier.

To visualize these results, we used a volcano plot, where points representing significant associations are colored by their direction of effect, and whether the species was identified as present in the sample by Metaphlan 4, Kraken2, or both (**Figure 4A-B**). We also represented this information in an UpSet plot, which visualizes the number of species that were found to be age associated in each cohort in both the individual tests and meta-analyzed results (**Figure 4C-D**). As a secondary visualization approach, we did not combine results across taxonomic methods, and instead we made volcano plots visualizing all age associations by method with lines connecting equivalent taxa (**Figure 4F-G**).
